# Epitope-based labeling for improved live-imaging of endogenous proteins in *C. elegans*

**DOI:** 10.64898/2026.02.05.703904

**Authors:** Elise van der Salm, Mette H. Schroeder, Loes B. Steller, Stephanie I. Miller, Amelie Scheper, Gwen Nowee, Erik. E. Griffin, Suzan Ruijtenberg

## Abstract

Visualizing protein expression dynamics with high temporal resolution is essential for understanding how cells acquire specific fates and functions during development, where key decisions can occur within minutes. Conventional direct fluorescent tagging often fails to capture these rapid changes in protein expression due to the relatively slow fluorophore maturation time. Indirect epitope-based labeling strategies offer a promising alternative, yet only a limited number of these systems have been developed and used in the context of multicellular organisms. Here, we evaluate and combine four epitope-based indirect labeling systems for live-imaging of proteins in *C. elegans*: the SunTag, Frankenbody, MoonTag and AlfaTag systems. Each system uses a fluorescently labeled high-affinity single-chain antibody or nanobody to recognize short peptide epitopes fused to a protein of interest, enabling immediate visualization of newly synthesized proteins. We demonstrate that all four systems specifically label epitope-tagged endogenous proteins and show no detectable cross-reactivity when used in dual-color combinations, enabling simultaneous visualization of distinct proteins within the same embryo. In addition, we show that the SunTag system offers three major advantages over direct labeling: earlier detection of proteins, enhanced sensitivity through signal amplification (as illustrated by CAM-1) and less impact on the function (as demonstrated for ERM-1). Together, this expanded toolkit of epitope-based labeling systems offers many new opportunities for visualizing rapid protein dynamics and for dissecting how their dynamics drive cell fate decisions during development.

**SUMMARY:** The development of epitope-labeling systems has improved live-imaging quality of proteins. Unfortunately, limited systems exist for multicellular organisms to study protein expression in the context of development. Here, we expand the epitope-labeling toolbox for *C. elegans* by combining SunTag or Frankenbody with MoonTag or AlfaTag. Our data indicates that these systems simultaneously visualize different endogenous proteins without cross-reactivity. Moreover, the SunTag system shows advantages over direct labeling: earlier detection, enhanced sensitivity through signal amplification and less impact on protein function. This expanded epitope-labeling toolbox in C. elegans provides opportunities for accurate visualization of different proteins that drive cell fate decisions.

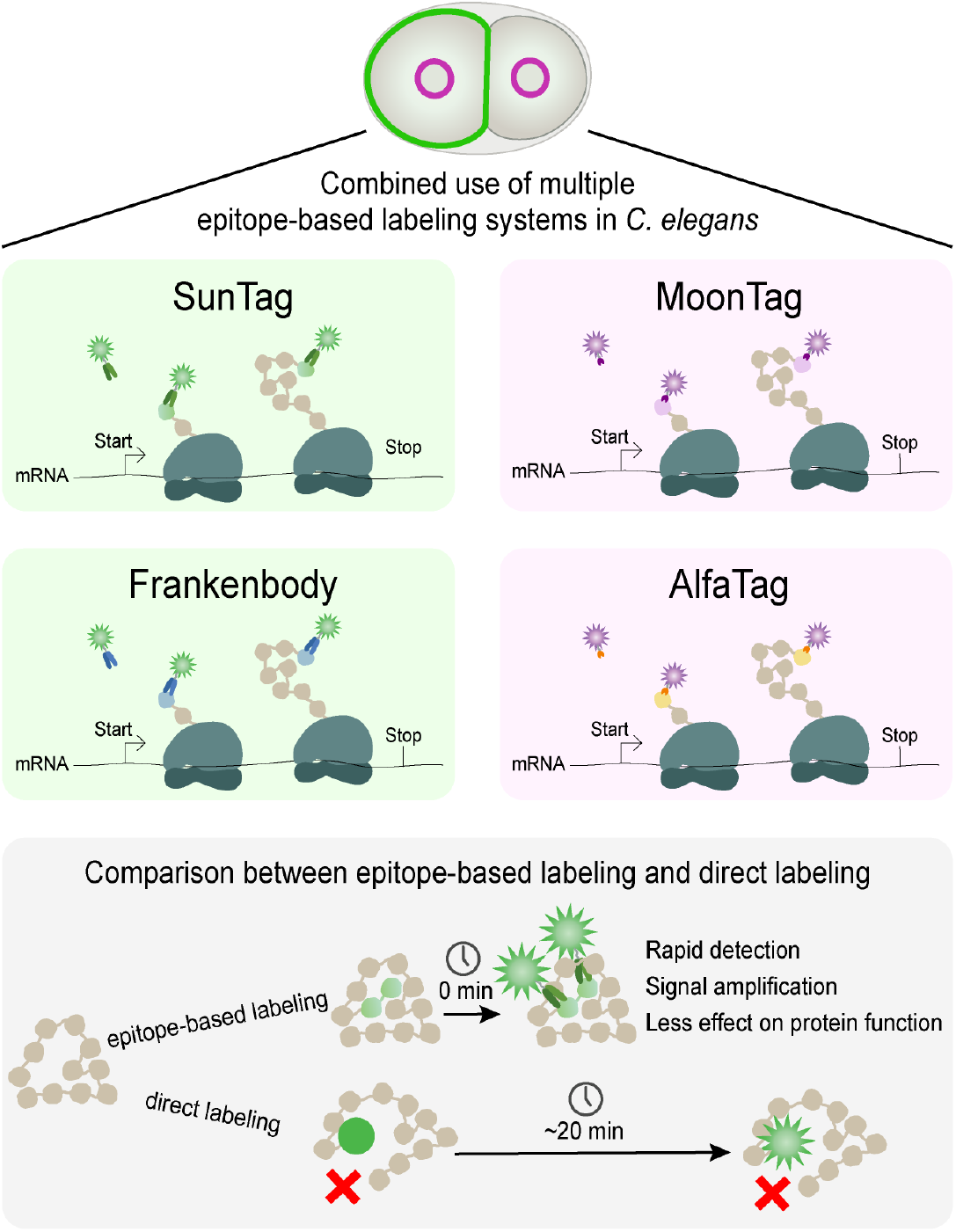

## Introduction

Precise regulation of protein expression is essential for development and homeostasis of multicellular organisms, including *Caenorhabditis elegans*. During development, cells make rapid fate decisions that depend on the abundance, location and activity of specific proteins. These changes in protein level can occur on timescales of minutes, and can differ across cell types even within the same organismal environment (Shukla et al. 2025; Sivaramakrishnan et al. 2023). Measuring and visualizing these rapid changes in protein abundance and activity is therefore essential for understanding the mechanisms underlying rapid cell fate acquisition during development. Capturing such dynamics requires live-imaging approaches with high temporal resolution that allow us to measure protein levels and subcellular localization. In most standard live-imaging approaches, a fluorescent protein is directly fused to the protein of interest (POI) to visualize its expression. However, direct fluorescent tagging often fails to detect rapid changes in expression because the maturation time of fluorescent proteins is often longer than the window of protein expression. As a result, newly synthesized proteins cannot be observed immediately after translation, and rapid and dynamic changes in protein expression may be missed (Shukla et al. 2025). In addition, the relatively large size of fluorescent proteins (∼25 kDa) can interfere with protein function or localization, making direct tagging unsuitable for some developmentally important proteins (Bergwell et al. 2019).

To overcome these limitations, several epitope-based labeling systems have been developed, including SunTag, Frankenbody, MoonTag and AlfaTag (Figure 1) (Akhuli et al. 2022; Boersma et al. 2019; Tanenbaum et al. 2014; Zhao et al. 2019). These indirect labeling systems use a single-chain variable antibody fragment (scFv) or nanobody fused to a fluorescent protein and that binds with high affinity to short epitopes fused to a POI. Because the fluorescently labeled scFv or nanobody is already matured before binding to the epitope, POIs become fluorescently labeled immediately upon synthesis of the fused epitope. This enables real-time visualization of nascent proteins and visualization of short-lived protein dynamics that cannot be observed with direct fluorescent tags. Furthermore, the small size of the epitopes minimizes perturbation of protein function and allows for signal amplification when multiple epitopes are incorporated (Bergwell et al. 2019; Virant et al. 2018). Moreover, incorporating an array of peptides allows long-term tracking of translation dynamics of single mRNAs and has been instrumental in many key discoveries in translation regulation (Morisaki et al. 2016; Pichon et al. 2016; Wang et al. 2016; Wu et al. 2016; Yan et al. 2016). Beyond live-imaging applications, epitope-based systems can be adapted for a wide range of purposes, as scFvs or nanobodies can be fused not only to fluorophores but also to various effector proteins. Such applications enable, for example, protein purification (Gotzke et al. 2019), transcriptional regulation at endogenous loci (Tanenbaum et al. 2014) and targeted DNA epigenome editing (Huang et al. 2017).

**Figure 1:**
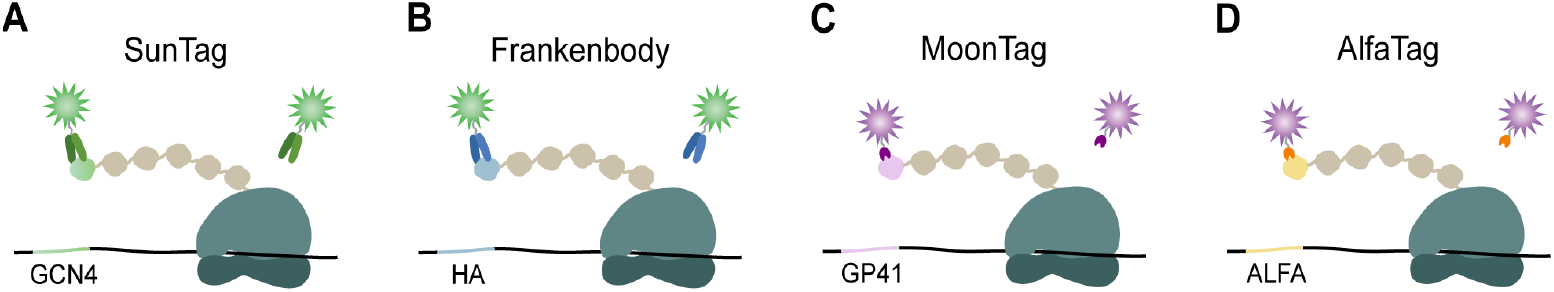
Schematic overview of the SunTag, Frankenbody, MoonTag and AlfaTag fluorescent protein labeling systems. **(A-D)** Schematics of four epitope-based labeling systems. SunTag and Frankenbody systems consist of a fluorescently labeled scFv that binds GCN4 **(A)** or HA **(B)** epitopes, respectively. MoonTag and AlfaTag systems consist of a fluorescently labeled nanobody that binds GP41 **(C)** or ALFA **(D)** epitopes, respectively. Binding occurs immediately upon synthesis of the epitope-tagged protein, enabling rapid visualization of the tagged protein of interest. In this study, SunTag and Frankenbody scFvs were fused to GFP (green), whereas MoonTag and AlfaTag were fused to HaloTag (purple), allowing combinations of systems for simultaneous labeling of multiple proteins.

Although epitope-based labeling systems are widely used in cell culture, only a limited subset has been adapted for use in multicellular model organisms (Boswell et al. 2025; Dufourt et al. 2021; Hu et al. 2023; Murakawa et al. 2022; Quintin et al. 2025; van der Salm et al. 2025a; van der Salm et al. 2025b; Vinter et al. 2021; Xu et al. 2022; Zhao et al. 2019). Yet, such tools are especially valuable in multicellular organisms, where gene expression and protein dynamics can change rapidly during development. In *C. elegans* for example, embryonic cells divide approximately every 20 minutes, requiring protein dynamics to be tightly regulated on similarly short timescales to ensure proper cell fate acquisition (Shukla et al. 2025; Sivaramakrishnan et al. 2023). Nevertheless, only SunTag and AlfaTag have been used in *C. elegans* for live-imaging of mRNA, translation or protein (Hu et al. 2023; Quintin et al. 2025; van der Salm et al. 2025a; van der Salm et al. 2025b). For most other systems, including the Frankenbody and MoonTag system, it is unclear whether they can be used in *C. elegans*. Moreover, in contrast to cellular systems (Vigano et al. 2021), it has not been tested whether multiple epitope-based labeling systems can be combined in *C. elegans* without cross-reactivity to visualize multiple proteins simultaneously. The limited development and systematic validation of epitope-based labeling strategies in multicellular model organisms constrain our understanding of complex biological processes in a developmental context.

Here, we evaluate four epitope-based labeling systems, SunTag, Frankenbody, MoonTag, and AlfaTag, for live-imaging of proteins in *C. elegans* embryos and larvae. All four systems use high-affinity scFvs or nanobodies to recognize short peptide epitopes. The SunTag and Frankenbody systems consist of a scFv that binds to either GCN4 or HA epitopes, respectively. The MoonTag and AlfaTag systems consist of a nanobody that binds with high affinity to either GP41 or ALFA epitopes, respectively (Figure 1). By combining GFP-labeled scFvs (named SunTag::GFP and Frankenbody::GFP throughout the paper) with Halo-labeled nanobodies (AlfaTag and MoonTag), we show that different proteins can be visualized simultaneously without detectable cross-reactivity. Furthermore, using the SunTag system we further demonstrate that epitope-based labeling systems enable more rapid detection of proteins and have a lower impact on protein function than direct fluorophore fusions. Together, these findings expand the toolkit of epitope-based labeling strategies for *C. elegans* and provide new opportunities for real-time visualization of rapid protein dynamics of multiple proteins during development.

## Results

### The SunTag, Frankenbody, MoonTag and AlfaTag systems label endogenous proteins without cross-reactivity

Recently, the SunTag and AlfaTag systems have been successfully adapted for protein labeling in *C. elegans*. We aimed to further expand the epitope-based labeling toolbox, and combine multiple systems for simultaneous visualization of different proteins. We focused on four systems: SunTag, Frankenbody, MoonTag and AlfaTag. In this project, both the SunTag and Frankenbody scFvs were labeled with GFP (SunTag::GFP and Frankenbody::GFP, respectively), and the MoonTag and AlfaTag nanobodies were fused to HaloTag (MoonTag::HALO and AlfaTag::HALO, respectively). HaloTags can be covalently bound to a range of fluorescent HaloTag ligands, providing spectral flexibility and facilitating combinations with other fluorophores within the same strain (Los et al. 2008). This design enables pairing either SunTag::GFP or Frankenbody::GFP with either MoonTag::HALO or AlfaTag::HALO. Each scFv or nanobody construct was integrated as a single-copy transgene into the *C. elegans* genome for stable expression. SunTag::GFP expression was driven by an *eft-3* promoter, expression of Frankenbody::GFP, MoonTag::HALO and AlfaTag::HALO was driven by a *mex-5* promotor. We first examined the distribution of SunTag::GFP, Frankenbody::GFP, MoonTag::HALO and AlfaTag::HALO in early *C. elegans* embryos in the absence of epitope-tagged target proteins. Under these conditions, all four systems showed uniform cytoplasmic localization with a slight enrichment in the nucleus (Figure 2A-D). Notably, SunTag::GFP and Frankenbody::GFP additionally showed enrichment in granule-like structures in the P-cell (Figures 2B and S1A) (Marnik et al. 2019). Next, we examined whether the SunTag, Frankenbody, MoonTag and AlfaTag systems can label endogenous proteins that are tagged with a single copy of their corresponding epitope. We also tested whether different systems can be combined to simultaneously visualize different proteins. To this end, we tested four different combinations: (1) SunTag with MoonTag, (2) SunTag with AlfaTag, (3) Frankenbody with MoonTag, and (4) Frankenbody with AlfaTag. The SunTag::GFP and Frankenbody::GFP were used to label PAR-3 (GCN4::PAR-3 and HA::PAR-3, respectively), which localizes to the cortex of the anterior cell AB at the 2-cell stage (Etemad-Moghadam et al. 1995). MoonTag::HALO and AlfaTag::HALO were used to label NPP-9, a nuclear pore complex protein (Nkombo Nkoula et al. 2023) (GP41::NPP-9 and ALFA::NPP-9, respectively) (Figure 2E-N). As expected, in all four combinations, MoonTag::HALO and AlfaTag::HALO localized to nuclear pores, while SunTag::GFP and Frankenbody::GFP were enriched at the cortex of the AB-cell. In addition, we observed some puncta in the P-cell, which could either be PAR-3 clusters or par-3 translation spots (Munro et al. 2004) (Figures 2F,G,K,L and S1B-C). Moreover, we did not observe enrichment of SunTag::GFP or Frankenbody::GFP at nuclear pores, nor enrichment of MoonTag::HALO or AlfaTag::HALO at the cell cortex, indicating that there is no cross-reactivity between the systems. Indeed, when we quantified fluorescent signals at the nuclear pore, we confirmed a strong and specific enrichment of MoonTag::HALO and AlfaTag::HALO (P ≤ 0.0001), while SunTag::GFP and Frankenbody::GFP remained at background levels (Figure 2H, I, M, N and S1D–G). Importantly, both SunTag::GFP and Frankenbody::GFP can robustly label NPP-9 if NPP-9 is fused to the corresponding GCN4 or HA epitopes, respectively (Figure S1H-L). Together, these results demonstrate that SunTag and Frankenbody can be combined with MoonTag and AlfaTag to simultaneously label different proteins without detectable cross-reactivity.

**Figure 2:**
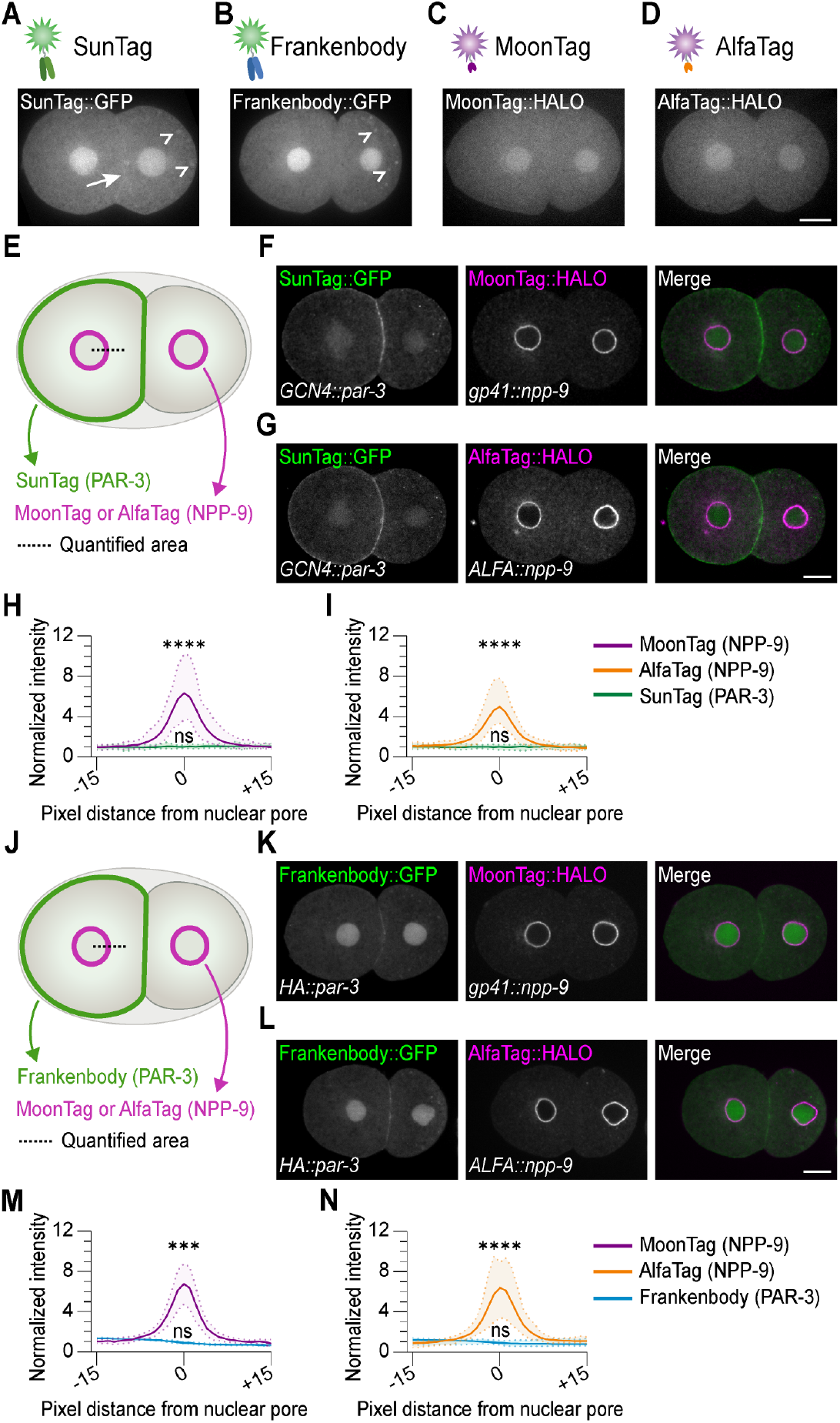
The SunTag, Frankenbody, MoonTag and AlfaTag systems label endogenous proteins in *C. elegans* embryos without detectable cross-reactivity. **(A-D)** Schematics of the four protein labeling systems and representative images of 2-cell stage C. elegans embryos expressing SunTag::GFP **(A)**, Frankenbody::GFP **(B)**, MoonTag::HALO **(C)**, or AlfaTag::HALO **(D)**. Arrow indicates enrichment at the membrane, open arrowheads indicate enrichment in granular structures. Scale bars: 10 μm. **(E)** Schematic of a 2-cell *C. elegans* embryo showing PAR-3 labeled with SunTag and NPP-9 labeled with either MoonTag or AlfaTag. **(F-G)** Representative images of 2-cell stage embryos expressing SunTag::GFP, GCN4::PAR-3 and either MoonTag::HALO and GP41::NPP-9 **(F)** or AlfaTag::HALO and ALFA::NPP-9 **(G)**. Images were acquired and displayed using identical settings. Scale bar: 10 μm. **(H-I)** Average fluorescence intensities of SunTag::GFP and MoonTag::HALO **(H)** or SunTag::GFP and AlfaTag::HALO **(I)** at the nuclear pore complex, as indicated in (E, dotted line), normalized to the mean nuclear and cytoplasmic intensity. Solid lines represent the mean and the shading dots represent the ± SD. Line scans (31 pixels long) were centered at pixel 0, marking the nuclear pore and extended to pixel −15 and +15 (*n*=27 embryos). Statistical test: A one-way ANOVA followed by a Bonferroni’s multiple comparison **(H)** or Kruskal–Wallis test followed by a Dunn’s multiple comparison test **(I)** comparing the background-corrected intensity at the nuclear pore (at pixel 0) to the mean nuclear and cytoplasmic intensity (at pixels −15 to −13 and +13 to +15). ns = not significant and ^****^P≤0.0001. **(J)** Schematic of a 2-cell stage *C. elegans* embryo showing PAR-3 labeled with Frankenbody::GFP and NPP-9 labeled with either MoonTag::HALO or AlfaTag::HALO. **(K-L)** Representative images of 2-cell stage embryos expressing Frankenbody::GFP, HA::PAR-3 and either MoonTag::HALO and GP41::NPP-9 **(K)** or AlfaTag::HALO and ALFA::NPP-9 **(L)**. Different brightness/contrast settings have been applied. Scale bar: 10 μm. **(M-N)** Average intensities of Frankenbody::GFP labeling together with MoonTag::HALO **(M)** or AlfaTag::HALO **(N)** as depicted in **(J)**, normalized to the mean nuclear and cytoplasmic intensity. Solid lines represent the mean and the shading dots represent the ± SD. Line scans (31 pixels long) were centered at pixel 0, marking the nuclear pore and extended to pixel −15 and +15 (*n*=18 embryos). Statistical test: A Kruskal–Wallis test followed by a Dunn’s multiple comparison test or one-way ANOVA followed by a Bonferroni’s multiple comparison test **(N)** comparing the background-corrected intensity at the nuclear pore (at pixel 0) to the mean nuclear and cytoplasmic intensity (at pixels −15 to −13 and +13 to +15). ns = not significant, ^***^P≤0.001 and ^****^P≤0.0001.

### SunTag-based labeling reveals newly synthesized proteins earlier than direct fluorescent tagging

Our next goal was to compare epitope-based labeling with direct protein labeling for visualization of newly synthesized proteins. A key advantage of epitope-based labeling is the immediate fluorescent labeling of nascent proteins due to the binding of already matured fluorophores. In comparison, direct fluorescent tagging requires fluorophore maturation, which can delay detection and may hinder correct interpretation of early protein localization and expression dynamics. Recently, Shukla et al. showed that although *cam-1* translation starts at the 2-cell stage, CAM-1 tagged with mNeonGreen (CAM-1::mNG) was not detectable by live-imaging until later stages. Importantly, fixation followed by incubation at 37 °C to allow fluorophore maturation revealed CAM-1 protein at the cell membranes of 4-cell stage embryos, indicating that delayed fluorophore maturation hindered early detection (Shukla et al. 2025). Based on these observations, we reasoned that CAM-1 provides an ideal test case to directly compare epitope-based labeling with conventional fluorescent protein tagging. To test this, we expressed SunTag::GFP together with CAM-1::2×GCN4 and observed clear enrichment of signal at the cell membranes at the 4-cell stage. This enrichment was not observed in embryos expressing SunTag::GFP alone or in embryos expressing CAM-1::mNG (Figure 3). Moreover, although CAM-1::mNG signal became detectable at later developmental stages (Figure 3F-G), the signal intensity was much lower than with SunTag-based labeling of CAM-1 (compare Figure 3E to 3F). Thus, SunTag labeling enables both earlier detection of endogenous proteins as well as signal amplification through multivalent epitope tagging.

**Figure 3:**
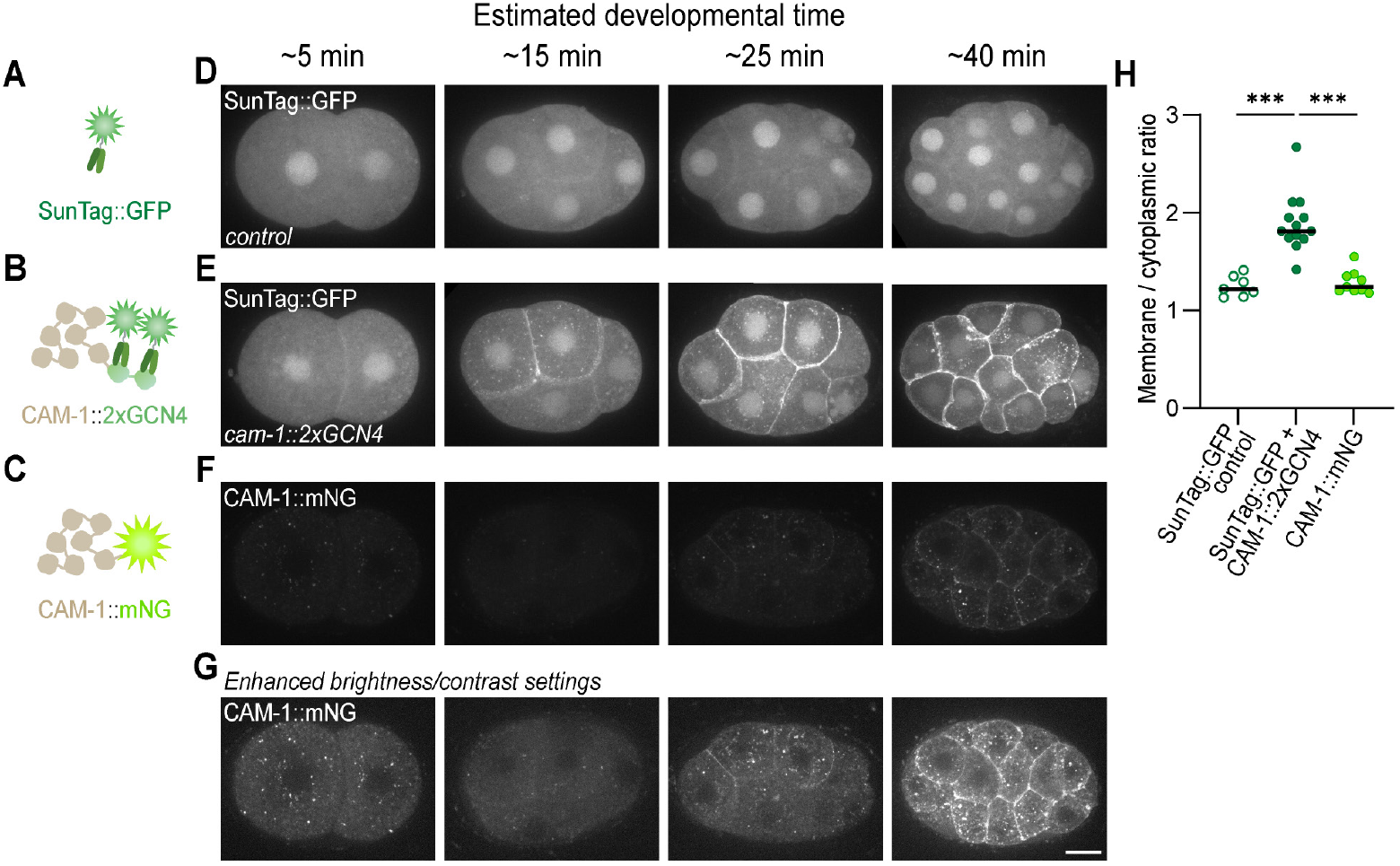
SunTag-based labeling reveals newly synthesized CAM-1 earlier than direct fluorescent tagging. **(A-C)** Schematic illustrations of the SunTag::GFP control **(A)**, SunTag::GFP-based labeling of CAM-1::2xGCN4 **(B)** and direct CAM-1 labeling using the conventional fluorophore mNeonGreen (CAM-1::mNG) **(C)**, representing the fluorescent labeling strategies in embryos shown in panels D-F. **(D-G)** Representative images showing the SunTag::GFP control **(D)** or CAM-1 localization at membranes using SunTag-based CAM-1 labeling **(E)** or direct CAM-1 labeling with CAM-1::mNG **(F)**, across developmental stages. Images in panels D-F were acquired and depicted using identical settings. Panel **(G)** shows CAM-1::mNG with enhanced brightness/contrast settings. Scale bar: 10 μm. **(H)** Quantifications of membrane-to-cytoplasmic ratio of CAM-1 levels for SunTag::GFP control (*n*=7), SunTag::GFP + CAM-1::2xGCN4 (*n*=13) and CAM-1::mNG (*n*=9). Each dot represents one embryo. Statistical test: Kruskal–Wallis test followed by a Dunn’s multiple comparison test. ^***^P≤0.001.

### SunTag labeling perturbs protein function less than labeling with conventional fluorescent proteins

Finally, we asked whether epitope-based labeling using the SunTag system preserves protein function better than direct labeling with conventional fluorescent proteins (Figure 4A-C). Fusion of relatively large fluorophores can interfere with protein activity, whereas smaller epitope tags are often better tolerated (Bergwell et al. 2019). To directly compare these approaches, we focused on ERM-1, a core cytoskeletal linker required for apical epithelial tissue integrity. ERM-1 function is highly sensitive to fluorescent tagging. Previous studies have reported that N-terminal fusion to ERM-1 disrupts protein function (Amieva et al. 1999; Henry et al. 1995; Roy et al. 1997), whereas C-terminal tagging, commonly used because it preserves viability, still causes defects in the excretory canal (Figure 4D), including canal extension defects and cyst formation (Ramalho et al. 2020). We first tested whether N-terminal tagging of ERM-1 is tolerated when using the SunTag system. In contrast to N-terminal GFP fusion, insertion of a single GCN4 epitope at the N-terminus of *erm-1a* resulted in viable animals expressing SunTag::GFP and GCN4::ERM-1a (abbreviated as SunTag::ERM-1a; Figure 4A,E). Thus, N-terminal tagging of ERM-1 is tolerated using small epitope tags. We next assessed ERM-1 functionality by analyzing the morphology of the excretory canal, a tissue that is highly sensitive to ERM-1 perturbation (Figure 4D-G). Despite clear expression of SunTag::ERM-1a in the excretory canals (Figure 4E), SunTag::ERM-1a animals displayed no detectable morphological defects (Figure 4H-I). It should be noted that because *erm-1a* and *erm-1b* differ at their N-termini, in this strain the ERM-1b isoform remains untagged and may contribute to canal formation. To tag both isoforms and directly compare functional consequences of indirect versus direct labeling, we examined strains in which both ERM-1 isoforms were tagged at the C-terminus, either using the SunTag system (expressing SunTag::GFP and ERM-1::GCN4; abbreviated as ERM-1::SunTag) (Figure 4B) or GFP (ERM-1::GFP) (Figure 4C) (Ramalho et al. 2020; van der Salm et al. 2025a). While ERM-1::GFP animals exhibited shortened excretory canals with frequent cysts and gaps, ERM-1::SunTag animals showed significantly milder defects and approximately 2.1-fold longer canals compared to ERM-1::GFP (Figure 4F-I). While we cannot exclude that not all ERM-1::GCN4 is bound by SunTag::GFP, it does indicate that the smaller tag is better tolerated compared to direct fluorophore tagging. Together, these results demonstrate that epitope-based labeling using the SunTag system interferes less with ERM-1 function than direct fluorescent protein fusion. More broadly, epitope-based approaches provide a powerful strategy for visualizing endogenous proteins while preserving function, enabling the study of dynamic protein localization and expression changes that may be inaccessible using conventional fluorescent tagging.

**Figure 4:**
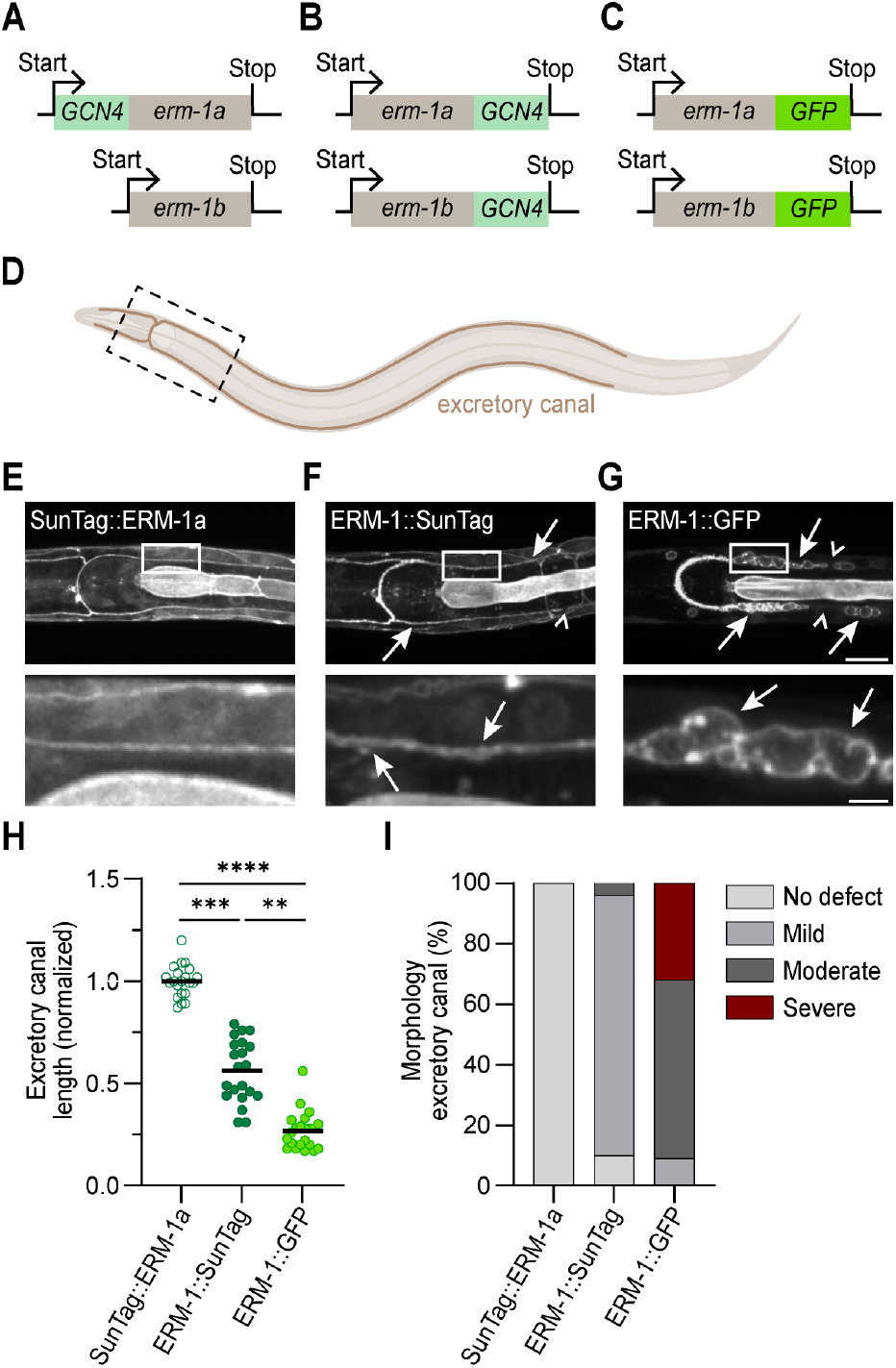
SunTag-based labeling perturbs protein function less than labeling with conventional fluorescent proteins. **(A-C)** Schematic illustrations of different ERM-1 fluorescent labeling strategies: N-terminal tagging of the ERM-1a isoform using 1xGCN4 (SunTag::ERM-1a; **A)**, and C-terminal tagging of both ERM-1 isoforms using 1xGCN4 (ERM-1::SunTag; **B)** or GFP (ERM-1::GFP; **C). (D)** Schematic illustration of the excretory canal in L1 C. elegans larvae, from a ventral view. The dashed box represents the depicted region in E-G. **(E-G)** Representative images showing the excretory canal of ERM-1::GFP **(E)**, ERM-1::SunTag **(F)** and SunTag::ERM-1a **(G)** L1 larvae from a ventral view. Insets below. Arrows indicate cysts, open arrowheads indicate gaps. Different brightness/contrast settings have been applied. **(H)** Quantification of the posterior excretory canal length of SunTag::ERM-1a, ERM-1::GFP and ERM-1::SunTag larvae (*n*=22, 21, 21). Data were normalized against the average excretory canal length of SunTag::ERM-1 larvae. Statistical test: Kruskal–Wallis test followed by a Dunn’s multiple comparison test. ^**^P≤0.01, ^***^P≤0.001, ^****^P≤0.0001. (**I**)Quantification of morphological defects in SunTag::ERM-1a, ERM-1::GFP and ERM-1::SunTag larvae (*n*=22, 21, 21).

## Discussion

In this study, we expanded the toolbox for epitope-based labeling for live-imaging in *C. elegans*. We examined four systems − SunTag, Frankenbody, MoonTag and AlfaTag − and tested whether different epitope-based labeling systems can be combined to visualize multiple proteins simultaneously. All tested systems specifically labeled endogenous proteins with high specificity and without cross-reactivity. These new developments in epitope-based labeling systems for the multicellular organism *C. elegans* broadens our possibilities for simultaneous live-imaging of endogenous proteins in the context of development.

### Epitope-based labeling versus direct protein labelling

Next to demonstrating specificity and combinatorial use of epitope-based labeling systems in *C. elegans*, our study highlights situations in which indirect labeling offers clear advantages over conventional direct fluorescent protein tagging. Direct labeling approaches require only a single fusion protein and do not rely on co-expression of an scFv or nanobody, making them experimentally straightforward. For proteins that are highly and stably expressed, direct fluorescent tagging can therefore be a convenient strategy for visualization and quantification. However, direct fluorescent labeling also has intrinsic limitations. Fluorophore maturation delays detection of newly synthesized proteins, which can hinder accurate interpretation of early protein localization and rapid expression dynamics. In addition, fusion of relatively large fluorescent proteins can interfere with protein folding or function, particularly for proteins that are structurally or functionally sensitive to tagging. Epitope-based labeling strategies can overcome several of these limitations. First, fusion of multiple epitopes enables signal amplification, allowing visualization of proteins that are expressed at very low levels and would otherwise remain undetectable with single fluorophores. Second, epitope-based labeling allows immediate detection of nascent proteins through binding of pre-matured fluorophores, enabling visualization across the full temporal window of protein expression. This is particularly important for proteins with rapid turnover, where relatively long fluorophore maturation times can limit detection of directly tagged proteins. Our CAM-1 experiments illustrate both of these advantages, as labeling CAM-1 with two GCN4 epitopes allowed earlier and brighter detection of CAM-1 compared to direct tagging with mNeonGreen. Third, the small size of epitope tags can reduce interference with protein folding and function, as demonstrated by our ERM-1 data where indirect SunTag labeling was better tolerated than direct GFP fusion. Taken together, epitope-based labeling systems are especially valuable to visualize proteins without the delay caused by fluorophore maturation, proteins that are lowly expressed or have fast turnover rates, and for proteins that are functionally sensitive to direct fluorophore tagging.

### Considerations for choosing an epitope-based labeling system

Our data demonstrate that all four tested epitope-based labeling systems are compatible with live-imaging in *C. elegans*. We did not observe one system to consistently outperform the others. However, practical considerations may influence which system is best suited for a given biological question. First, when expressed in the absence of epitope-tagged target proteins, all four systems − SunTag, Frankenbody, MoonTag, and AlfaTag − showed diffuse cytoplasmic localization with a slight enrichment in the nucleus. This baseline nuclear enrichment is particularly important to consider when labeling nuclear proteins, such as transcription factors, as nuclear protein accumulation may be masked by high background levels of scFv or nanobody in the nucleus. In addition, we observed system-specific localization patterns that may interfere with certain experiments. The Frankenbody::GFP scFv displayed enrichment in P-granules, which could complicate its use for visualizing proteins that localize to, or in close proximity to, P-granules. We previously reported that the SunTag::GFP scFv exhibits off-target affinity for epithelial apical junctions at later developmental stages, although this did not interfere with its ability to bind epitope-tagged proteins (van der Salm et al. 2025a). Interestingly, we observed a similar off-target localization for the MoonTag system when fused to GFP and expressed broadly in somatic tissues (strain not used in this study). Because Frankenbody::GFP and AlfaTag::HALO expression was driven by the *mex-5* promoter, which is restricted to the germline and early embryos, we were unable to assess whether these systems also display apical junction affinity once epithelial tissues are established. Such off-target localization may need to be considered when studying membrane-associated proteins in differentiated tissues.

### Adaptations to fine-tune live-imaging of proteins

Building upon these system-specific features, our results highlight several practical considerations for optimizing epitope-based labeling in *C. elegans*. All indirect labeling strategies require a careful balance in the relative expression levels of the epitope-tagged target protein and the unbound fluorescent scFv/nanobody, as this balance determines the signal-to-noise ratio and the visibility of the labeled target protein. If scFv/nanobody expression is too low, not all epitope-tagged proteins will be labeled, whereas too high expression increases background fluorescence from unbound scFv/nanobodies, masking specific signal of the target protein. Optimal scFv/ nanobody levels therefore depend on the expression of the target protein, with highly expressed proteins requiring higher scFv/nanobody levels than low-abundance targets. In this study, we used different promoters to control scFv/ nanobody expression levels. SunTag::GFP scFv was expressed under control of the *eft-3* promoter, which drives broad expression in all tissues, but results in relatively low expression in embryos. In contrast, Frankenbody::GFP, MoonTag::HALO, and AlfaTag::HALO constructs were expressed under the *mex-5* promoter, which results in high expression in the germline and early embryos. While both promoters enabled robust live-imaging, lower scFv expression seemed to be advantageous for visualizing proteins expressed at low levels, as for example PAR-3. Membrane localization of PAR-3 was clearer using the *eft-3*-driven SunTag::GFP system than with the *mex-5*-driven Frankenbody::GFP system (Figure 2F–G versus 2K–L). This difference is not because of reduced binding of the Frankenbody::GFP, as peak membrane intensities were comparable or slightly higher in the Frankenbody system, but instead resulted from higher background in the *mex-5* driven system, which reduced the signal-to-background ratio and masked membrane localization. These results indicate that promoter selection can be used to tune signal-to-noise ratio and optimize visualization of proteins across a wide range of expression levels.

Fluorophore choice represents an additional layer of optimization. To date, SunTag scFvs binding GCN4 have primarily been fused to GFP, which improves solubility and minimizes aggregation (Tanenbaum et al. 2014). However, alternative fluorophores may provide advantages in brightness, photostability, or spectral flexibility depending on experimental needs. In our study, MoonTag and AlfaTag nanobodies were fused to HaloTag, enabling covalent labeling with different ligands and facilitating multicolor imaging. Although the HaloTag-based strategy provides sufficient signal for localization studies, variability in ligand uptake into the embryo between experiments and individual animals suggests that in many animals, not all MoonTag::HALO or AlfaTag::HALO were fluorescently labeled, limiting reliable quantification of protein levels. In addition, ligand staining increases experimental complexity and may vary across tissues or developmental stages. These limitations could potentially be overcome by replacing HaloTag with directly fused fluorescent proteins, as for example AlfaTag has previously been fused to mKate in *C. elegans* (Quintin et al. 2025), and we have successfully fused MoonTag to GFP (data not shown).

## Conclusion

In summary, we established the SunTag, Frankenbody, MoonTag and AlfaTag systems as useful tools for epitope-based labeling of endogenous proteins in *C. elegans*. By evaluating specificity and cross-reactivity, we demonstrate that multiple systems can be used simultaneously to visualize multiple different proteins in a developmental context. Compared to direct fluorophore tagging, these epitope-based systems allow earlier detection and improved sensitivity as illustrated by our CAM-1 data. Moreover, they interfere less with protein function, as demonstrated for ERM-1, making them particularly valuable for proteins with rapid turnover or for targets whose function is disrupted by the addition of large fluorescent proteins. At the same time, our results highlight the importance of optimizing scFv or nanobody expression levels to optimize signal-to-noise ratios for specific proteins or experiments. Together, this expanded epitope-based labeling toolkit for *C. elegans* provides new opportunities to visualize multiple endogenous proteins with high temporal resolution during development. By enabling simultaneous imaging of protein localization and dynamics, these approaches open the door to studying how coordinated changes in protein abundance and localization drive cell fate decisions and tissue patterning.

## Acknowledgments

We thank L.S. Sackmann for help with experiments. We also thank Ruijtenberg lab members, S. van den Heuvel, M. Boxem, S. Suijkerbuijk and L. Braccioli for helpful discussions. We thank S. van den Heuvel for support with funding. This work utilized resources from Utrecht University. We also thank WormBase and the Biology Imaging Center Faculty of Sciences of the Department of Biology at Utrecht University. Some strains were provided by the Caenorhabditis Genetics Center, which is funded by the US National Institutes of Health (NIH) Office of Research Infrastructure Programs (P40 OD010440). We thank M. Boxem for the BOX213 strain and D.L. Updike for the pgl-1(sam52[pgl-1::mTAGRFP::3XFLAG]) IV strain.

## Data availability

Data are available upon request.

## Supplemental information

Figures S1

Table S1. Excel file containing additional data too large to fit in a PDF, related to Methods

Table S2. Excel file containing additional data too large to fit in a PDF, related to Methods

## Methods

### *C. elegans* strains and maintenance

All strain names and associated genotypes can be found in Table S1. Worm strains were cultured following standard procedures (Brenner 1974). Worms were grown on nematode growth medium (NGM) agar plates seeded with OP50 *Escherichia coli* (*E. coli*) at 15 °C or 20 °C.

### Strain generation

All *C. elegans* strains described in this study were generated using CRISPR/Cas9 for single-copy integration of transgenes; a detailed overview of all transgenes, including used repair templates and sgRNAs/crRNAs is provided in Table S2. For single-copy integrations at MosSCI loci, a plasmid-based CRISPR/Cas9 approach was used (Friedland et al. 2013). Injection mixes containing eft-3p::Cas9 plasmid (50 ng/μL; Addgene plasmid #46168), sgRNA plasmid (50 ng/μL), repair template plasmid (50 ng/μL) and co-injection marker plasmid (2.5 ng/μL) were delivered by microinjection into the gonads of young adult animals. Offspring expressing the co-injection marker were grown, lysed and genotyped to select for genome-edited animals. All other single-copy insertions were performed using a Cas9 ribonucleoprotein-based method (Ghanta et al. 2021). Injection mixes contained Cas9 protein (0.25 μg/μL) (IDT), TracRNA (0.1 μg/μL) (IDT), pRF4 [rol-6(su1006)] (40 ng/μL) co-injection marker (Mello et al. 1991), locus-specific crRNA (56 ng/μL) (IDT) and either ssODN (0.11 μg/μL) (IDT) or melted dsDNA (25 ng/μL) repair template, or 100 ng/uL of plasmid pSM63 (GFP::Frankenbody repair template). dsDNA repair templates were synthesized by PCR using 5’ SP9 modified primers (IDT) and purified using the NucleoSpin Gel and PCR Clean-up Kit (Macherey-Nagel, 740609.250). For strains generated by crossing, hermaphrodites were heat-shocked at 30 °C for 6 h and recovered at 20 °C for 3 days to obtain males. Males were crossed with relevant strains and progeny was genotyped.

### HaloTag labeling

HaloTag labeling was performed as previously described, by culturing animals in S-medium provided with OP50 and halo ligand (Borchers et al. 2025; Chang and Dickinson 2022). In short, to prepare S-medium, autoclaved S-basal [NaCl (5.844 g/L), K2HPO4^*^3H2O (11.4115 g/L) and KH2PO4 (6.8045 g/L)] was provided with 1% (vol/vol) autoclaved trace-metal solution [disodium EDTA (5 mM), FeSO5 (2.5 mM), MnCl2 (1 mM), ZnSO4 (1 mM) and CuSO4 (0.1 mM)], 1% (vol/vol) Potassium Citrate (1 M, pH 6.0), 0.3% (vol/vol) MgSO4 (1 M) and 0.3% (vol/vol) CaCl2+ (1M). The medium was filter-sterilized before adding 0.1% (vol/vol) cholesterol (5 mg/mL in EtOH), 1% (vol/vol) Penicillin-Streptomycin (P4458, Sigma) and 0.1% Nystatin Suspension (N1638, Sigma Aldrich). For each labeling reaction, 30 L4 hermaphrodites were incubated in 30 μL S-medium containing 10 μL OP50 *E. coli* pellet and 0.375 μL Halo ligand (200 μM stock) in an Eppendorf tube at 20 °C and 150 rpm. After overnight incubation, animals were transferred to NGM agar plates with OP50 and allowed to recover for 1-3 h. SUR113 and SUR114 animals were labeled using JFX646 HaloTag ligand (Lavis Lab), SUR219 and SUR220 were labeled using JFX650 HaloTag ligand (Lavis Lab).

### Microscopy and Image Analysis

Gravid adults were splayed in M9 buffer [KH2PO4 (0.22 M), Na2HPO4 (0.42 M), NaCl (0.85 M) and MgSO4 (0.001 M)] with or without tetramisole (10 mM) to release embryos. Embryos were mounted on a 5% agarose pad. Spinning-disk confocal imaging was performed on an Eclipse Ti2-E with perfect focus spinning disk (Nikon, Kōnan Japan) equipped with a CSU-X1-M1 confocal head (Yokogawa, Tokio Japan), Prime BSI sCMOS camera and CFI Plan Apo λ 100×/1.45 oil or CFI Plan Apo λ 60x/1.4 oil objective (see Table S1). Identical imaging settings were used within experiments, unless stated otherwise. Images were acquired using MetaMorph Microscopy Automation and image analysis software. Images were analyzed and processed using FIJI (Schindelin et al. 2012).

### Quantification of PAR-3 labeling

To examine PAR-3 enrichment, 2-cell embryos were imaged with 0.25 μm intervals along the z-axis. Maximum intensity projections were generated from 5 selected z-stacks for each embryo using FIJI (Schindelin et al. 2012). PAR-3 labeling was quantified from a 3-pixel-wide line scan, drawn along the membrane between the AB and P-cell. Line scan intensities were averaged before subtracting mean background intensities measured from a region outside the embryo. Mean cytoplasmic values were measured from a region in the AB-cell and background corrected. To normalize data, background-corrected PAR-3 labeling values were divided by background-corrected cytoplasmic values.

### Quantification of NPP-9 intensity and cross-reactivity

To assess NPP-9 labeling, 2-cell and 4-cell stage embryos were imaged with 0.25 μm intervals along the z-axis and analyzed using FIJI (Schindelin et al. 2012). For each embryo, a maximum intensity projection was generated from 5 selected z-stacks encompassing the nuclear envelope. Fluorescent signal at the nuclear pore was quantified from a 1-pixel-wide, 31-pixel long line scan, drawn centered at and perpendicular to the nuclear membrane. To examine cross-reactivity, the position of the line scan was defined based on the HaloTag fluorescence, which was the only channel showing detectable nuclear pore signal, and used for quantification of both channels. Average background fluorescence was measured from a region outside the embryo and subtracted from the line scan intensities. Fluorescence intensities were normalized against unbound scFv::GFP or nanobody::HALO signal in the nucleus and cytoplasm, which was calculated from the average of pixel values −15 to −13 and +13 to +15.

### Quantification of CAM-1

Embryos expressing CAM-1::mNG or SunTag::GFP, with or without co-expression of CAM-1::2xGCN4, were imaged at the 4-cell stage with 0.4 μm z-step increments. FIJI (Schindelin et al. 2012) was used to generate maximum intensity projections of 3 selected z-stacks encompassing the ABa-ABp membrane. Membrane-associated CAM-1 signal was quantified from the peak histogram intensity of a 100 pixel wide line scan drawn perpendicular to the ABa-ABp membrane. Cytoplasmic CAM-1 intensity was measured from a region within the ABa cell and averaged. Background fluorescence was measured from a region outside the embryo and averaged. After background subtraction, membrane CAM-1 levels were normalized to cytoplasmic CAM-1 levels to obtain the membrane-to-cytoplasm ratio.

### Quantification of excretory canal length and morphology

For excretory canal length and morphology analysis, adults were bleached to obtain synchronized L1 larvae (5 hours post hatching) expressing ERM-1::GFP, or SunTag::GFP combined with 1xGCN4::ERM-1a or ERM-1::1xGCN4. L1 larvae were imaged with 0.4 μm z-step increments. Of each animal, the length of the longest posterior excretory canal was measured using the Freehand Line tool in FIJI (Schindelin et al. 2012). The morphology of the excretory canals was assessed in the same individuals and scored as ‘no defects’ (no irregularities), ‘mild defects’ (small cysts), ‘moderate defects’ (larger cysts and/or little gaps) or ‘severe defects’ (severe cysts and/or frequent large gaps).

### Statistical analysis

Statistical analyses were performed using GraphPad Prism v.10. Data distributions were assessed for normality using a D’Agostino-Pearson test. For normally distributed data, comparisons of two populations were performed using an unpaired Student’s t-test, and comparisons of more than two populations were performed using a one-way ANOVA followed by a Bonferroni’s multiple comparison test. For data not drawn from a normal distribution, comparisons of two populations were made using a Mann–Whitney test, and comparison of more than two populations were carried out using a Kruskal–Wallis test followed by a Dunn’s multiple comparison test. NPP-9 labeling (represented in line-scan figures) was analyzed using background-normalized intensity values, to compare nuclear pore intensities at pixel 0 with the average intensities of unbound scFv::GFP or nanobody::HALO at pixel −15 to −13 and +13 to +15.

## Supplementary information

**Figure S1:**
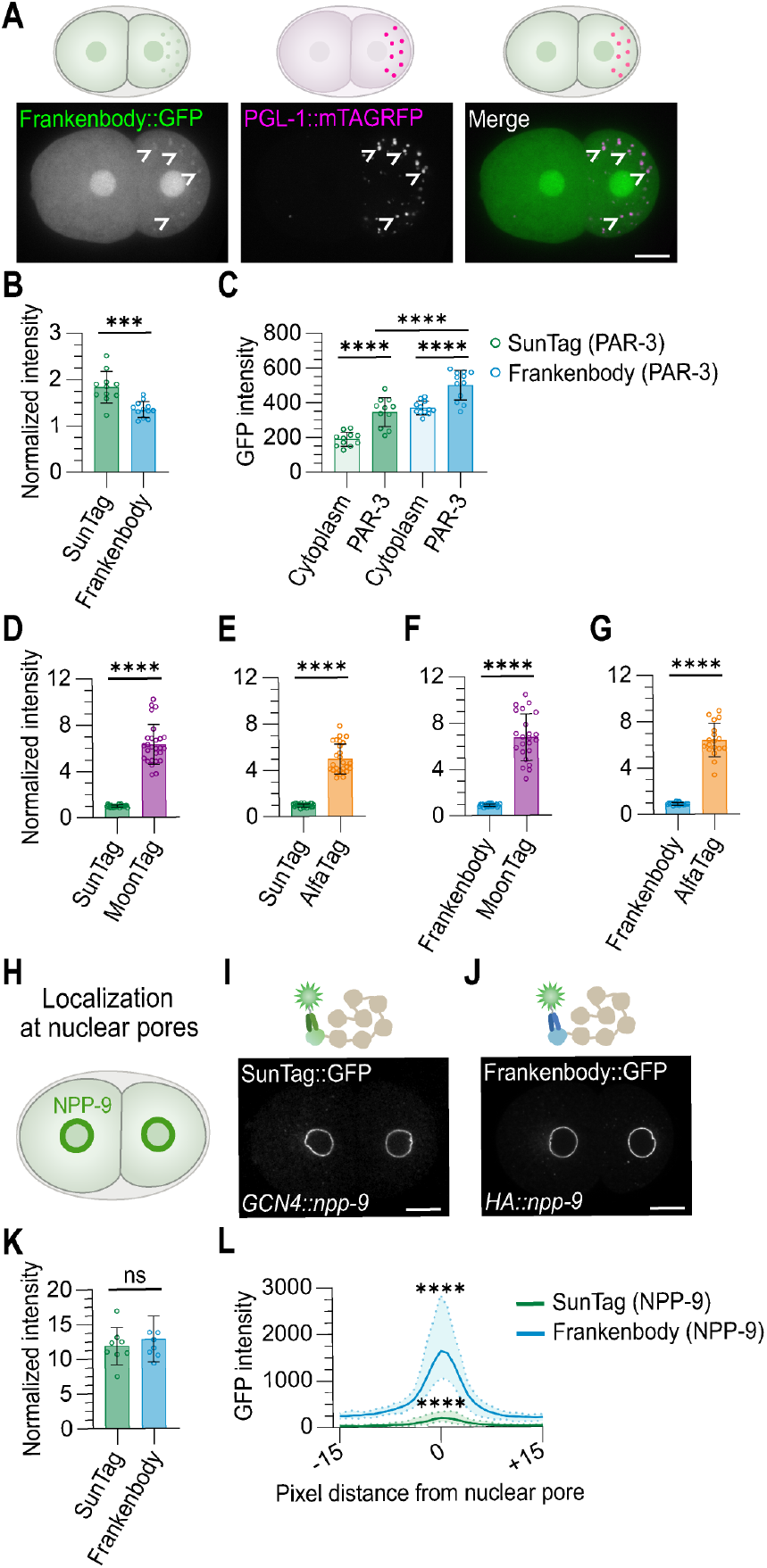
Characteristics of different epitope-labeling systems (related to Figure 2). **(A)** Schematic and representative images of a 2-cell C. elegans embryo expressing Frankenbody::GFP and the P granule marker PGL-1::mTAGRFP. Open arrowheads indicate P granules. Scale bar: 10 μm. **(B)** Relative PAR-3 fluorescence at the cell membrane between the AB and P-cell of embryos expressing either SunTag::GFP and GCN4::PAR-3, or Frankenbody::GFP and HA::PAR-3 (*n*=11 for SunTag, *n*=12 for Frankenbody). Statistical test: Unpaired Student’s t-test. ^***^P≤0.001. **(C)** Background-corrected fluorescence of PAR-3 in the cytoplasm and at the cell membrane between the AB and P-cell in embryos expressing either SunTag::GFP and GCN4::PAR-3, or Frankenbody::GFP and HA::PAR-3 (*n*=11 for SunTag, *n*=12 for Frankenbody). Data has been re-used from Figure S1B. Statistical test: one-way ANOVA followed by a Bonferroni’s multiple comparison test. ^***^P≤0.001. **(D-G)** Relative fluorescence intensities at the nuclear pore of embryos expressing SunTag::GFP, GCN4::PAR-3, MoonTag::HALO and GP41::NPP-9 **(D)**, SunTag::GFP, GCN4::PAR-3, AlfaTag::HALO and ALFA::NPP-9 **(E)**, Frankenbody::GFP, HA::PAR-3, MoonTag::HALO and GP41::NPP-9 **(F)** or Frankenbody::GFP, HA::PAR-3, AlfaTag::HALO and ALFA::NPP-9 **(G)**, normalized to the mean nuclear and cytoplasmic intensity. Quantifications are based on pixel value 0 in Figure 2H,I,M,N. Statistical test: two-tailed unpaired Student’s t-test. ^****^P≤0.0001. **(H)** Schematic illustration of SunTag::GFP or Frankenbody::GFP labeling endogenous NPP-9 at nuclear pore complexes. **(I-J)** Representative images of NPP-9 labeling in embryos expressing SunTag::GFP and GCN4::NPP-9 **(I)** or Frankenbody::GFP and HA::NPP-9 **(J)**. Different brightness/ contrast settings have been applied. Scale bars: 10 μm. **(K)** Relative fluorescence intensities at the nuclear pore of embryos expressing either SunTag::GFP and GCN4::NPP-9, or Frankenbody::GFP and HA::NPP-9 (*n*=8 embryos for each condition). Statistical test: Mann–Whitney test. ^***^P≤0.001 **(L)** Average fluorescence intensities of embryos expressing either SunTag::GFP and GCN4::NPP-9, or Frankenbody::GFP and HA::NPP-9 at the nuclear pore complex, normalized to the mean nuclear and cytoplasmic intensity. Solid lines represent the mean and the shading dots represent the ± SD. Line scans (31 pixels long) were centered at pixel 0, marking the nuclear pore and extended to pixel −15 and +15 (*n*=8 embryos for each condition). Data has been re-used from Figure S1G. Statistical test: Unpaired Student’s t-tests, performed on each dataset separately, comparing the background-corrected intensity at the nuclear pore (at pixel 0) to the mean nuclear and cytoplasmic intensity (at pixels −15 to −13 and +13 to +15). ^****^P≤0.0001.

